# Meta-analysis of the brain transcriptomes of multiple genetic mouse models of schizophrenia highlights dysregulation in striatum and thalamus

**DOI:** 10.1101/2025.03.26.645496

**Authors:** Kira A. Perzel Mandell, Sean K. Simmons, Ajay Nadig, Zohreh Farsi, Wei-Chao Huang, Sameer Aryal, Min Jee Kwon, Bryan Song, Kira Brenner, Nate Shepard, Ally A. Nicolella, Lesley D. O’Brien, Aron H. Lichtman, Joshua Z. Levin, Morgan Sheng

## Abstract

Schizophrenia is a severe mental illness with high heritability, but its underlying mechanisms are poorly understood. We meta-analyzed large-scale brain transcriptomic data from mice harboring individual loss-of-function mutations in seven schizophrenia risk genes (*Akap11*, *Dagla*, *Gria3*, *Grin2a*, *Sp4*, *Srrm2*, *Zmym2*). While all studied brain regions were affected, the striatum and the thalamus emerged as key brain regions of convergence. Striatum showed downregulation of synapse-and oxidative phosphorylation-related gene sets in all models. In the thalamus, mutants separated into two groups based on transcriptomic phenotype: synapse-related gene sets were upregulated in mutants with only schizophrenia and bipolar association, and were downregulated in mutants that are associated with developmental delay/intellectual disability in addition to schizophrenia. Overall, our meta-analysis reveals convergence and divergence in brain transcriptomic phenotype in these schizophrenia genetic models, supports the involvement of striatal disturbance and synapse dysfunction in schizophrenia, and points to a key role of the thalamus.

## INTRODUCTION

Schizophrenia (SCZ) is a severe mental illness that affects approximately 0.7% of people globally ^1^. Despite being highly heritable, with an estimated heritability of around 80% ^2^, its underlying pathophysiology is inadequately understood. Genetic studies have identified many associations with schizophrenia, with the latest GWAS identifying 287 significantly associated loci ^3^. However, these common variant associations have low effect sizes (odds ratio typically < 1.2) and are usually in non-coding regions of the genome ^4^.

A recent study from the Schizophrenia Exome Sequencing Meta-Analysis (SCHEMA) consortium identified rare coding variants which confer substantial risk for schizophrenia, implicating 10 genes at an exome-wide significance level, and 32 genes at a false discovery rate (FDR) < 5% ^5^. These “SCHEMA genes” (a term we use here to include the FDR < 0.05 genes) harbor protein-truncating variants (PTVs) and damaging missense variants that were found to be associated with schizophrenia. Unlike common variants, these variants have a large effect on disease risk (odds ratio 2-50) and clearly cause loss of function of a specific gene, making them better suited for biological investigation. The study of these SCHEMA genes may allow us to gain insight into the functional molecular changes underlying schizophrenia by understanding the function of the genes themselves and also by understanding the consequences of their loss-of-function (LoF) in animal models ^6^.

The schizophrenia risk genes identified in the SCHEMA study have diverse molecular functions. In this study we focus on seven of these genes for which we have extensive RNA-seq data from mutant mice. Three of them have functions directly related to synaptic signaling.

*Grin2a* encodes a subunit of the N-methyl-D-aspartate (NMDA) receptor, a postsynaptic glutamate receptor that is implicated in schizophrenia by psychopharmacology ^7^, and is also a common-variant risk gene by GWAS fine-mapping ^3^. Mice heterozygous for a LoF mutation of *Grin2a* model several aspects of schizophrenia and show large-scale transcriptomic changes across the brain ^8^. *Gria3* encodes an AMPA receptor subunit - another subtype of postsynaptic glutamate receptor ^9^. *Dagla* encodes diacylglycerol lipase alpha, an enzyme involved in the biosynthesis of the endocannabinoid, 2-arachidonoylglycerol (2-AG) that plays a role in synaptic signaling ^10^. Three other selected SCHEMA genes have functions related to transcription and mRNA processing. *Sp4*, which encodes a transcription factor, has additionally been genetically linked to schizophrenia through GWAS fine-mapping ^3^, and may also have human genetic association with bipolar disorder (BP) ^11^. *Srrm2* encodes an RNA binding protein, localized in nuclear speckles and involved in RNA splicing ^12^. *Zmym2* encodes a zinc-finger protein which has been implicated in transcriptional regulation through its binding and modulation of transcription factors and histone modifiers ^13,14^. Finally, *Akap11* encodes an A-kinase anchoring protein, which interacts with several signaling factors including PKA ^15^. *Akap11* LoF mutations are also associated with bipolar disorder ^16^.

As with many complex diseases, schizophrenia has overlapping genetic risk with other disorders, such as bipolar disorder as noted above ^17^. A sizable fraction of genes identified in the SCHEMA study (SCHEMA genes) also show an association with neurodevelopmental disorders, developmental delay, and intellectual disability (DD/ID) ^5,16^. Rare coding variants in *Srrm2* and *Zmym2* showed an association with DD/ID when tested in a separate, previously published DD/ID cohort ^18^. On the other hand, *Akap11*, *Sp4*, *Gria3*, and *Dagla* do not have known associations to DD/ID and appear to be relatively “SCZ/BP-specific.” *Grin2a* shows genetic association with DD/ID, however there is allelic segregation-schizophrenia is associated with protein-truncating LoF variants whereas DD/ID is associated with missense mutations including dominant-negative gain-of-function mutations ^5,19^.

While SCHEMA genes are diverse in their molecular functions and disease associations, they have in common their association to schizophrenia. Thus, we hypothesized that mouse mutants in SCHEMA genes would show some degree of phenotypic convergence in the brain - as measured by systematic transcriptomic analysis (bulk RNA-seq) of multiple brain regions - which might give insights into the molecular mechanisms and systems-level brain alterations of schizophrenia.

## RESULTS

### Widespread changes in brain transcriptome among SCHEMA mouse mutants

In order to understand the effect of schizophrenia risk gene heterozygous loss of function in the brain, knockout mouse lines were studied for seven individual genes identified by SCHEMA. These genes were *Gria3* ^20^, *Grin2a* ^8^, *Sp4* ^21^, *Akap11* ^22^, *Dagla*, *Srrm2* ^23^, and *Zmym2* ^24^. As a comparator, we also included a mouse mutant that carries a null mutation (C456Y) in *Grin2b* ^8,25^, a gene that is not associated with schizophrenia, but is associated with autism spectrum disorder ^26^ (ASD) and DD/ID ^27^.

We focused our meta-analysis on heterozygous mutant mice because heterozygosity is the genotype of SCHEMA cases. Furthermore, for *Srrm2*, *Zmym2*, and *Grin2b*, the homozygous mutant is not viable, and for *Gria3*, an X-linked gene, the mutant male mice had a hemizygous genotype (-/y). Bulk RNA sequencing was performed to measure the transcriptomic changes caused by these SCHEMA gene mutations (compared with wild-type litter-mate controls), examining six brain regions (hippocampus, prefrontal cortex (PFC), substantia nigra, striatum, somatosensory cortex (SSC), and thalamus) at 1 and 3 months of age (Supplemental Figure 1A, Supplemental Table 1).

Based on the number of differentially expressed genes (DEGs, defined as differentially expressed with FDR < 0.05, see Methods, Supplemental Table 2), all tested brain regions and ages were affected by these SCHEMA mutations (+/-or-/y). By DEG count, the most highly affected brain region and the most affected age varied across SCHEMA mutants (Figure 1A). For example, the *Sp4* mutant had more DEGs at 1 month than 3 months and had the greatest number of DEGs in the substantia nigra, whereas *Grin2a* overall had more DEGs at 3 months and the most DEGs in the PFC. *Sp4*, *Srrm2*, and *Zmym2* mutants in general showed more DEGs than the other SCHEMA mutants (Figure 1A), which is perhaps not surprising since *Sp4*, *Srrm2*, and *Zmym2* are known to have direct functions related to transcription and RNA processing. The *Sp4* analysis also included more biological replicates which could increase the number of DEGs detected. *Grin2a* and *Grin2b* mutants had many DEGs but interestingly their trends over age differed - *Grin2a* had more DEGs at 3 months, while *Grin2b* had more DEGs at 1 month, correlating with the earlier expression of *Grin2b* and the later expression of *Grin2a* during brain development in mice ^28^.

**Figure 1:**
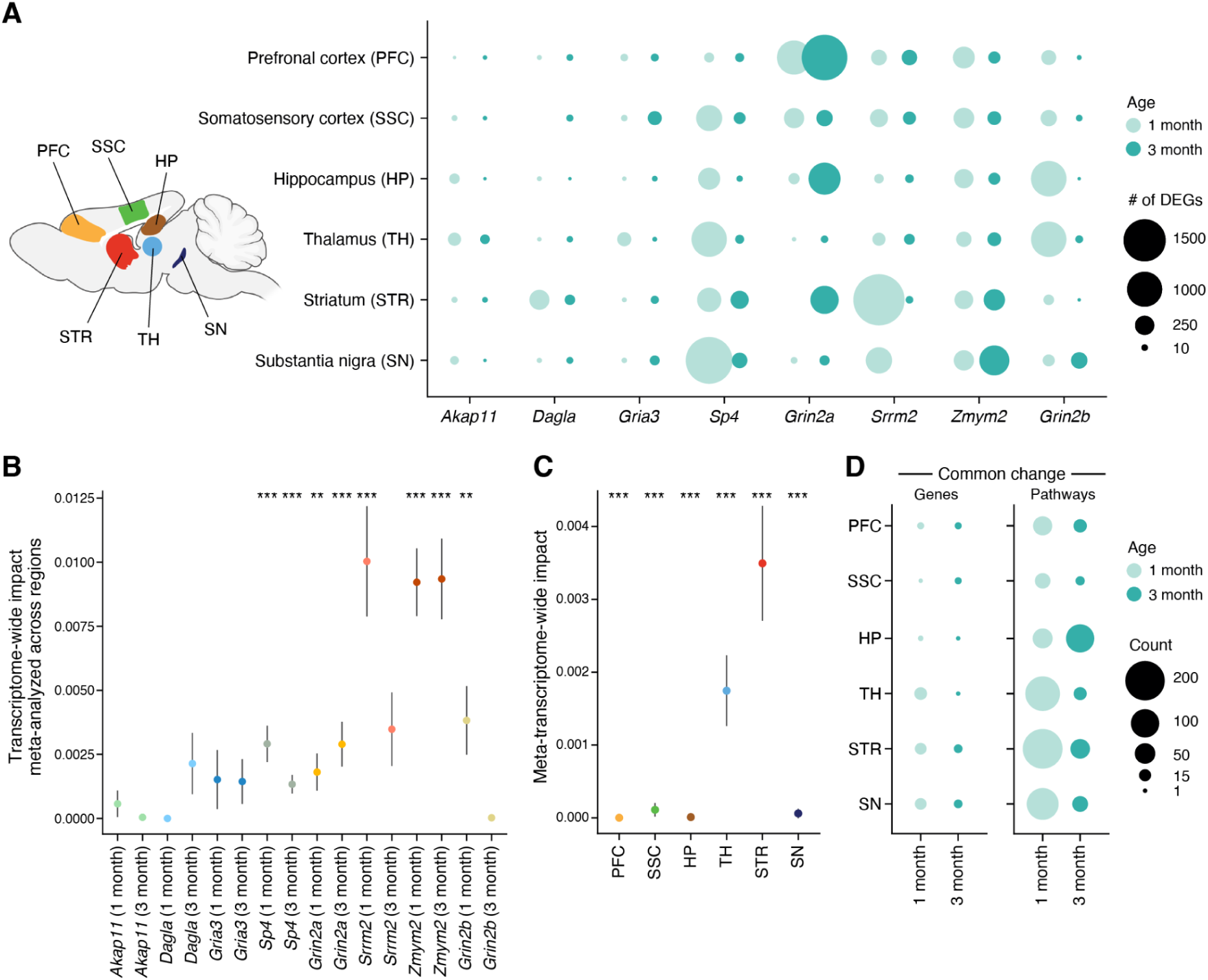
Overview of transcriptomic impact of SCHEMA mutants and Grin2b. **A)** Number of differentially expressed genes (DEGs) for each mutant, brain region, and age. DEGs defined by FDR < 0.05. Color represents age and size represents number of DEGs. **B)** Transcriptome wide impact (TWI) of each mutant at each age. TWI was meta-analyzed across all brain regions within a mutant and age. Error bars represent estimated standard error (see Methods). **C)** Meta-transcriptome wide impact in each brain region. TWI was meta-analyzed across SCHEMA mutants and ages for each brain region. **D)** Number of overlapping genes and pathways across SCHEMA mutants for each brain region and age. Overlapping as defined by p < 0.05 in ≥ 4 SCHEMA mutants. Stars indicate significance, *: FDR < 0.05, **: FDR < 0.01, ***: FDR < 0.001

While counting significant DEGs can provide insight into magnitude of transcriptomic change, this measure is biased by statistical power, such that better-powered experiments (for instance, those with greater n number) will tend to yield more significant DEGs. To mitigate this potential source of bias, we additionally performed Transcriptome-Wide Analysis of Differential Expression (TRADE, see Methods) ^29^. TRADE estimates the distribution of differential expression effects across the whole transcriptome, rather than individual genes, allowing us to measure transcriptome-wide impact while avoiding bias from sample size. We estimated the transcriptome-wide impact (TWI) for each individual brain region at each age for each SCHEMA mouse mutant (Supplemental Figure 1B, Supplemental Table 3), and we also meta-analyzed the TWI scores for each mutant across brain regions to assess broader patterns across different mutants (Figure 1B, Supplemental Table 3, see Methods). This TRADE analysis corroborated that mutants in SCHEMA genes with transcriptional or nuclear function (*Sp4, Srrm2, Zmym2*) had generally larger effects on the transcriptome than others such as *Akap11* and *Dagla* (Figure 1B, Supplemental Figure 1B). TRADE analysis also confirmed that different mutants show different patterns in transcriptome-wide impact between 1 and 3 months. For example, *Sp4* and *Srrm2* and *Grin2b* had a greater TWI at 1 month than at 3 months, while *Grin2a* and *Dagla* had a greater TWI at 3 months. *Zmym2* had a consistently strong TWI across both ages (Figure 1B). Thus DEG count and TWI score are largely consistent with each other (compare Figure 1A and Supplemental Figure 1B).

### Striatum and thalamus show the greatest and most convergent transcriptomic impact across SCHEMA mutants

A key systems-level question that we wanted to address is, which brain regions are particularly affected in SCHEMA mutant mice? Meta-analysis of transcriptome-wide impact scores by brain region across ages and SCHEMA mouse mutants showed that the striatum had the highest meta-TWI score, followed by thalamus (Figure 1C, Supplemental Table 3). The other regions (PFC, SSC, HP, SN) had much lower meta-TWI scores–although all are statistically significant, owing in part to the large number of samples meta-analyzed (Figure 1C). At this level of statistical power, even very small effects can be significant, so in this analysis, the effect size is more important than the p-value.

Notably, when this meta-analysis was separated by age, different age dynamics can be observed in different brain regions. The PFC had a greater meta-TWI at 3 months than at 1 month, whereas the SSC and thalamus had a greater meta-TWI at 1 month than at 3 months (Supplemental Figure 1C). This is interesting because the PFC, a brain region much implicated in SCZ pathophysiology ^30^, is thought to mature at a later age than other parts of the cortex ^31,32^.

Besides assessing the magnitude of overall transcriptomic impact, we also aimed to identify any specific gene expression changes that were common across many SCHEMA mutants. At the individual gene level, in each brain region at each age, we searched for genes that were altered (nominal p < 0.05) in the same direction in at least half (≥4) of the SCHEMA mutants. Although there were a relatively small number of such “common-change” genes that fit the above criteria, they were found across the brain and were most abundant in the striatum, thalamus, and substantia nigra, and generally more at 1 month than 3 months of age (Figure 1D, Supplemental Table 4).

To explore transcriptomic changes at the “pathway” level, we performed gene set enrichment analysis (GSEA) on the differential expression results (see Methods, Supplemental Table 5). To identify pathway changes shared across mutants, we again searched for Gene Ontology (GO) terms that were changed with nominal significance (p < 0.05) in the same direction in at least half of the SCHEMA mutants for each of the specific brain regions and ages. There were considerably more such common-change pathways than common-change genes (Figure 1D, Supplemental Table 4), suggesting that gene set enrichment may be a helpful avenue to identify convergences across mutant models. With the exception of the hippocampus, there were more common-change pathways at 1 month than at 3 months of age in the various brain regions. Based on the number of common-change pathways, the striatum at 1 month was the brain region that showed the greatest overlap across the studied SCHEMA mutants, followed by thalamus and substantia nigra (Figure 1D). This was partially related to the total number of significant GSEA terms across all mutants in each region. For example, at 1 month, hippocampus, PFC, and SSC had fewer total significantly affected pathways across mutants (1557-2459), while striatum, thalamus, and substantia nigra had more (3129-4594). However, this relationship was not linear, because striatum had the highest number of common-change pathways across SCHEMA mutants despite having fewer total changed pathways than thalamus and substantia nigra (Supplemental Figure 1D, Supplemental Table 2). This supports the hypothesis that the highest level of biological convergence across SCHEMA mutants occurs in the striatum.

We also performed joint analysis of the different mouse models in specific brain regions and at specific ages by performing Stouffer’s Z-score meta-analysis at both the gene and pathway level (Supplemental Table 4,6). At 1 month, striatum, thalamus, and substantia nigra again stood out as having the highest numbers of significantly altered genes and pathways (Supplemental Figure 1E). There were overall fewer significant genes and pathways at 3 months, but at the pathway level, striatum and substantia nigra still had relatively high numbers, along with hippocampus. Together with the transcriptome-wide impact scores, these findings indicate that striatum and thalamus are the brain regions most strongly impacted in SCHEMA mouse mutants; and furthermore, striatum and thalamus show the greatest similarity of transcriptomic change across SCHEMA mutants, closely followed by substantia nigra, particularly at the 1 month age.

### SCHEMA mutant phenotypes converge on several pathways in the striatum (synapse, mitochondrial respiration)

The striatum has been implicated in schizophrenia ^33^, not least because it receives major dopaminergic input and expresses high levels of D2 dopamine receptors - the target of most antipsychotic medications ^34^. As discussed above, in a systematic analysis of multiple brain regions, we found that the striatum showed the highest transcriptome-wide impact and high overlap of transcriptomic change across SCHEMA mutant mouse models, particularly at 1 month of age (Figure 1). To assess which pathways are commonly changed in the striatum across the different SCHEMA mutants, we performed a meta-analysis on GO GSEA results from the 1 month striatum (Figure 2A). This revealed 873 significant GO terms that were altered between SCHEMA mutants vs. their wild-type controls (FDR < 0.05, Supplemental Table 6).

**Figure 2:**
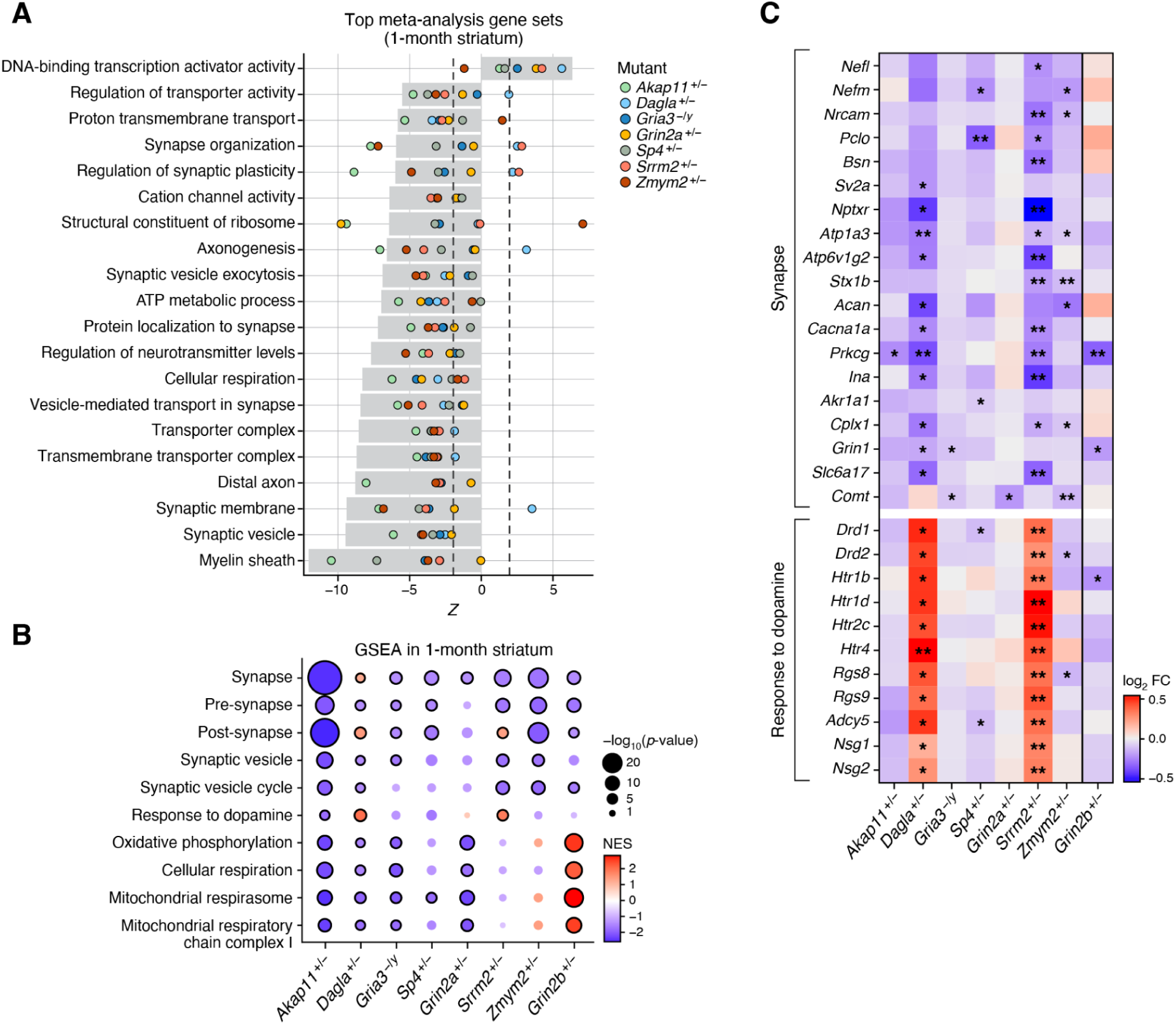
Convergent changes across SCHEMA mutants in 1 month Striatum. **A)** Top 20 most significant gene set changes by meta-analysis in 1 month striatum across SCHEMA mutants. Gray bar represents Stouffer’s meta Z score, each point represents the individual gene set Z scores from each SCHEMA mutant. Values outside the dashed lines correspond to the Z score which represents nominal significance (p < 0.05). For the purpose of display, redundant gene sets were collapsed and only the strongest of the similar gene sets is plotted (see Methods). **B)** Individual GSEA results for top convergent synapse-and OXPHOS-related gene sets. Color represents normalized enrichment score (NES), with red indicating upregulation and blue indicating downregulation. Bubble size represents significance, and black borders indicate FDR < 0.05 significance. **C)** Synapse-related genes which are consistently downregulated across SCHEMA mutants and dopamine response-related genes which are consistently upregulated in Srrm2 and Dagla mutants. Synapse related genes were selected from rank-sum analysis and Stouffer’s meta-analysis to prioritize genes which were downregulated across all SCHEMA mutants. All listed “synapse” genes are significantly (FDR < 0.05) downregulated in meta-analysis. *: p < 0.05, **: FDR < 0.05.

Many of the top gene sets from meta-analysis of striatum were related to synapses (Figure 2A-B), showing a general downregulation of synapse-related terms across SCHEMA mouse mutants as well as in the *Grin2b* mutant (Figure 2B, Supplemental Table 5). Among the many synapse-related gene sets, terms related to the presynapse and synaptic vesicle (e.g. “synaptic vesicle”, “vesicle-mediated transport in synapses”) were the most consistently downregulated across SCHEMA mutants. Other gene sets related more generally to synapses (such as synaptic membrane, synapse organization, synaptic plasticity) or to postsynapse were also significantly altered across SCHEMA mutants, mostly in a downward direction, with the exception of *Dagla* and *Srrm2* mutants, in which some of the synapse-related gene sets were upregulated (Figure 2A-B, Supplemental Figure 2A).

The individual genes underlying these synaptic pathway changes were variable between mutants, but a number of mRNA changes were largely directionally consistent (Figure 2C).

Meta-analysis of the gene-level differential expression results from each experiment showed that many synaptic or synaptic transmission-related genes were significantly downregulated across mutants, including Piccolo (*Pclo*), *Sv2a*, Bassoon (*Bsn*), and syntaxin 1B (*Stx1b*)(Figure 2C, Supplemental Table 7), which are related to the presynapse and synaptic vesicle dynamics. Notably, *Pclo* ^35^ and *Sv2a* ^36^ have growing evidence of genetic association with schizophrenia. Other downregulated genes in the broad “synapse” GO term include neurofilament light chain (*Nefl*) ^37^, neurofilament heavy chain (*Nefm*), and neuronal pentraxin receptor (*Nptxr*)^38^, whose protein products have been identified as potential biomarkers of neuronal injury and neurodegeneration. *Grin1* is the main essential subunit of the NMDA receptor, and its downregulation across SCHEMA mutants is consistent with the NMDA hypofunction hypothesis of schizophrenia ^39^ (Figure 2C, Supplemental Table 2).

Gene sets related to cellular respiration and oxidative phosphorylation, such as “cellular respiration”, “mitochondrial respiratory chain complex I”, and “mitochondrial respirasome” were significantly affected in striatum (Figure 2A-B), suggesting altered brain energetic metabolism in SCHEMA mutants. Interestingly, while these mitochondrial respiration gene sets showed downregulation in striatum across the studied SCHEMA mutants, they were upregulated in the striatum of the *Grin2b* mutant (Figure 2B). This distinguishing feature might reflect a difference in striatal dysfunction between mouse mutants of genes associated with schizophrenia versus DD/ID. In fact, hyperactivation of mitochondrial respiration complexes has been previously reported in the striatum of a *Fmr1*-knockout mouse model of the neurodevelopmental disorder Fragile X syndrome^40^. While mitochondrial respiration-related gene sets were altered in many brain regions in different mutants, the specific pattern of these pathways showing opposite and significant change in SCHEMA mutants (SCZ) versus *Grin2b* mutant (ASD) was found only in striatum (Figure 2B). At the gene level, the differential expression effect sizes of oxidative phosphorylation genes in *Akap11* and *Grin2a* mutants (which show the strongest downregulation of the gene set) showed significant anti-correlation with the changes in the *Grin2b* mutant (Supplemental Figure 2B). Of further note, the downregulation in mitochondrial respiration-related gene sets in striatum is strongest in the SCHEMA mutants with no PTV association with DD/ID, whereas the signal is weaker or absent in *Srrm2* and *Zmym2*, the SCHEMA mutants that show an association with DD/ID and, in this respect, are more similar to *Grin2b* (Figure 2B). We speculate that the divergence of transcriptomic phenotype with respect to mitochondrial respiration in the striatum may reflect pathophysiological network differences between genetic mouse models of schizophrenia versus DD/ID and ASD. However, characterization of additional mouse mutants representing these classes will be needed to validate this hypothesis.

Overall *Srrm2* and *Dagla* mutants showed especially high correlation in their transcriptomic effects in 1 month striatum, despite their apparently very different molecular functions (Supplemental Figure 2C). Of particular interest, the *Srrm2* and *Dagla* mutants had a significant (FDR < 0.05) upregulation of the “response to dopamine” gene set in the striatum, including increased expression of *Drd1* and *Drd2* (Figure 2B-C), which are dopamine receptors highly relevant to schizophrenia ^33^. Besides dopamine receptors, *Srrm2* and *Dagla* mutants also showed upregulation of several serotonin receptor genes (*Htr1b*, *Htr1d*, *Htr2c*, *Htr4*) as well as genes involved in GPCR signaling that could play roles downstream of dopamine and serotonin receptors (*Adcy5*, *Rgs8*, *Rgs9*) (Figure 2C). A reduction of *Comt*, which is a dopamine-metabolizing enzyme, has been demonstrated at both the RNA and protein level in the brains of *Grin2a* heterozygous and homozygous mutants ^8^. Here we find that *Comt* mRNA also drops in 1 month striatum of *Gria3* and *Zmym2* mouse mutants (Figure 2C). However, at 3 months, with the exception of the *Grin2a* mutant, this trend of *Comt* downregulation is not consistent (Supplemental Table 2).

The most statistically significant of the GO meta-analysis changes in striatum was a downregulation of the “myelin sheath” gene set across SCHEMA mutants (Figure 2A). However, while the “myelin sheath” gene set was significantly downregulated across five of the SCHEMA mutants, GO terms more specifically related to myelin, such as “myelination” and “ensheathment of neurons”, were significantly downregulated only in *Akap11* and *Sp4* mutants (Supplemental Figure 2D). This apparent discrepancy suggests that the “myelin sheath” finding is partially driven by genes not directly related to myelination; indeed, the most consistently decreased genes in the “myelin sheath” gene set across SCHEMA mutants were largely related to glycolysis (ex: *Gapdh, Aldoa, Pkm, Pgam1*) and the cytoskeleton (ex: *Nefm, Nefl, Plec, Tuba1b, Tubb4b*)(Supplemental Figure 2E).

At 3 months of age, the extent of pathway convergence in striatum across SCHEMA mutant models (as measured by number of significant pathways in Stouffer’s meta-analysis) was more limited than at 1 month, showing fewer and more weakly significant gene sets from GSEA meta-analysis (Supplemental Figure 1E, 2F). For instance, synapse-and cellular respiration-related gene sets were no longer significantly downregulated in meta-analysis across SCHEMA mutants at 3 months in striatum. In some individual SCHEMA models these pathways were downregulated, but in others they were upregulated or little-changed.

### Heterogeneity of transcriptomic changes in the PFC across SCHEMA mouse mutants

The PFC is highly developed in humans and primates and PFC dysfunction is implicated in schizophrenia ^41,42^. The PFC showed nominally significant transcriptome-wide impact scores at 1 or 3 months of age for all SCHEMA mutants, with the exception of *Akap11* and *Dagla*, for which the TWI score did not reach statistical significance in any brain region at 1 or 3 months of age (Supplemental Figure 1B, Supplemental Table 3). Significant TWI in this many SCHEMA mutants did not occur in any other brain region. Interestingly, the *Grin2b* heterozygous mutant did not show a significant TWI in the PFC, despite having significant impact on most other brain regions. This supports a recent report suggesting that the PFC is not as perturbed as other cortical regions in ASD ^43^. The extent of transcriptome-wide impact on the PFC notwithstanding, the PFC showed lower levels of convergence across the SCHEMA mouse mutants compared to other brain regions (Figure 1D, Supplemental Figure 1E). Our meta-analysis showed that the specific gene-and pathway-level changes in PFC were quite heterogeneous across the different SCHEMA mutant models, with at best modest correlations between SCHEMA mutants’ differential expression (such as *Sp4* and *Grin2a*; R = 0.31; Supplemental Figure 3A). In fact, the two SCHEMA genes with the most similar molecular function, *Gria3* and *Grin2a* - both encoding glutamate receptor subunits - were notably anti-correlated in their effects on the PFC transcriptome (R =-0.46; Supplemental Figure 3A).

At the 1 month time point, some of the top hits from GSEA meta-analysis of PFC included the downregulation of myelin sheath, cellular respiration, presynapse-and synaptic vesicle-related terms (Figure 3A, Supplemental Table 6), which is similar to the striatum (Figure 2A). However, these terms in PFC were not as consistently or robustly downregulated across all mutants as in striatum, and in most cases, the GSEA result reached FDR-significance at the individual level for only a few SCHEMA mutants and sometimes changed in opposite directions (Figure 3B). Furthermore, *Grin2b* mutants showed changes similar to those in SCHEMA mice in many of these pathways in PFC (Figure 3B, Supplemental Table 5). Overall, synapse-related gene sets were commonly changed in the PFC in SCHEMA mutants, but they were less directionally consistent or less frequently significant than in the striatum, which is in line with the lower number of common-gene or common-pathway changes in PFC (see Figure 1D).

**Figure 3:**
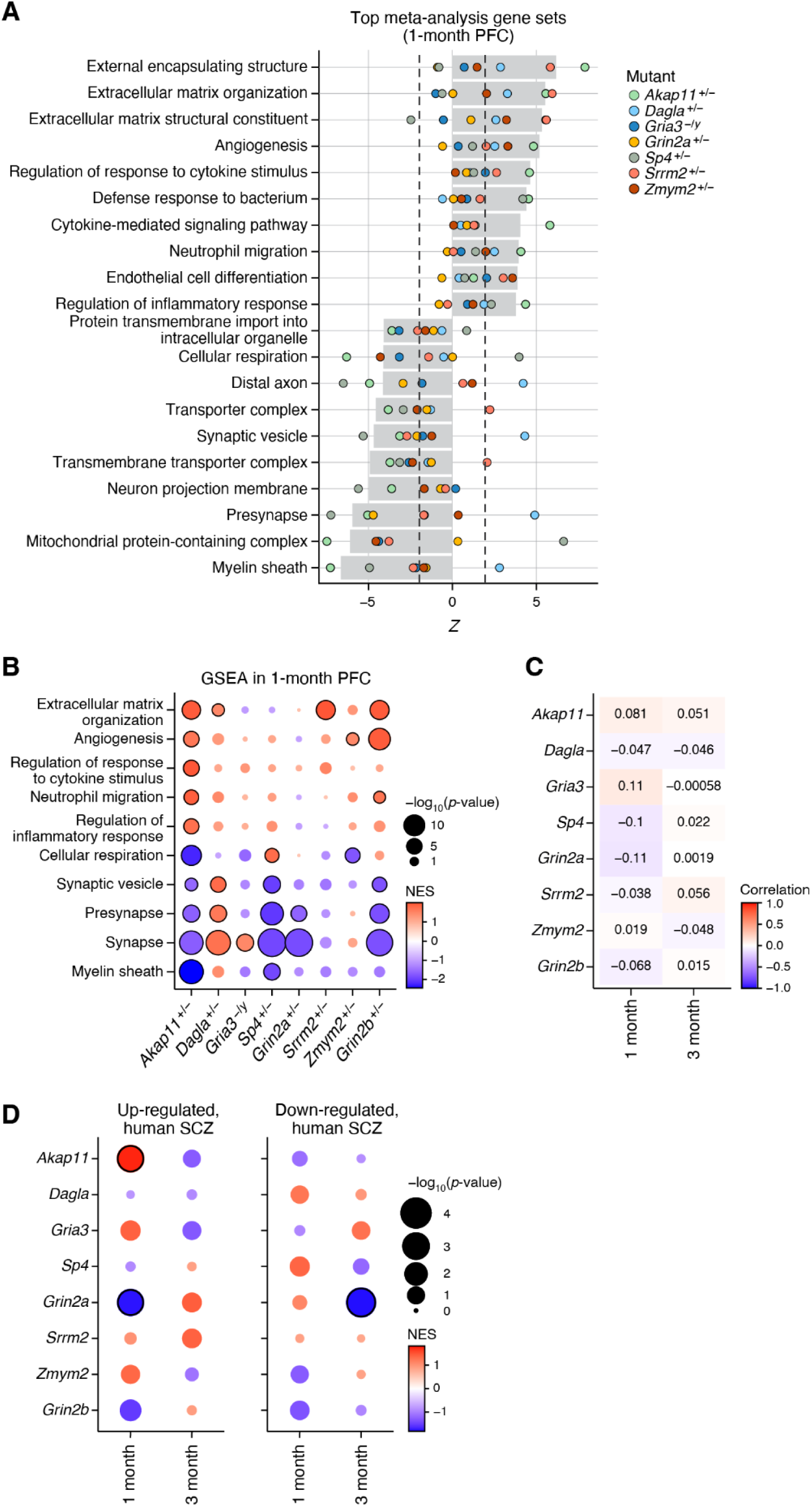
Greater heterogeneity of transcriptomic change in the PFC across SCHEMA mutants. **A)** Top 20 most significant gene set changes by meta-analysis in 1 month PFC across SCHEMA mutants. Gray bar represents Stouffer’s meta Z score, each point represents the individual gene set Z scores from each SCHEMA mutant. Values outside the dashed lines correspond to the Z score which represents nominal significance (p < 0.05). For the purpose of display, redundant gene sets were collapsed and only the strongest of the similar gene sets is plotted (see Methods). **B)** Individual GSEA results from top meta-analysis and synapse gene sets. Color represents normalized enrichment score (NES), with red indicating upregulation and blue indicating downregulation. Bubble size represents significance, and black borders indicate FDR < 0.05 significance. **C)** Correlation of differential expression results from bulk RNA sequencing between SCHEMA mutant PFC and human SCZ postmortem PFC. Correlation was performed on the union of nominally significant genes from each pair of results. Color and number indicate R value for each mutant and age. **D)** GSEA for upregulated DEGs and downregulated DEGs identified in human patient postmortem PFC.

Like 1 month PFC and 1 month striatum, “myelin sheath” was also the most significantly changed term in 3 month old PFC (Supplemental Figure 3B), being significantly downregulated in the *Akap11*, *Dagla*, *Grin2a*, and *Sp4* mutants. However, like in 1 month striatum, the “myelination” gene set is only significantly downregulated in the *Akap11* and *Sp4* mutants (Supplemental Figure 3C). The most consistently downregulated genes in this gene set across all SCHEMA mutants were more related to cytoskeleton terms (ex: *Actb, Actg1, Arf6, Tuba1b*), axons (ex: *Nefh, Dpysl2*), and mitochondria (ex: *Hspa9, Immt, Prdx3, Cox5a*) rather than the process of myelination itself (Supplemental Figure 3D).

Gene expression differences in human PFC have been studied by RNA-seq using schizophrenic and non-schizophrenic post-mortem brain tissues, so we compared our differential expression results to those from one of the largest studies of this kind ^44^. Perhaps unsurprisingly since the human brain samples are highly heterogeneous, whereas our brain samples are derived from mice of defined sex, age and similar genetic background, the bulk transcriptomic results were overall weakly correlated, if at all, between SCHEMA mutant mice PFC and human schizophrenic PFC (Figure 3C, see Methods). We created gene sets of the upregulated DEGs and downregulated DEGs from the human study ^44^ and performed GSEA using these annotated gene sets. Among the SCHEMA mutants studied, only *Grin2a* and *Akap11* had significant enrichment of human DEGs in their up-or down-regulated genes.

Notably, downregulated DEGs from human study were enriched among downregulated genes in the PFC of *Grin2a* mutants at 3 months, the age at which *Grin2a* mutants show the most transcriptomic changes. In the case of *Akap11* mutant, upregulated DEGs in human SCZ were significantly enriched among upregulated genes in the PFC at 1 month (Figure 3D, Supplemental Table 8). We note that *Akap11* mutant brain also showed some similarity to human SCZ PFC in proteomics studies of synapses ^42^.

### Thalamus shows a convergence within non-DD/ID-associated mutants and divergence with DD/ID-associated mutants

Though the thalamus transcriptome is less studied than PFC or striatum in relation to schizophrenia, we found a high degree of transcriptome-wide impact and a high level of transcriptomic convergence across SCHEMA models in the thalamus, especially at 1 month of age (Figure 1C-D). Meta-analysis of the thalamus GSEA results showed that the majority of the most significantly altered gene sets were upregulated, in contrast to the striatum in which the majority of altered gene sets were downregulated (compare Figure 4A with 2A). As observed for striatum, the gene sets with the highest Z scores in thalamus were synapse-related – here we observed upregulation of both presynaptic and postsynaptic terms (Supplemental Table 6).

**Figure 4:**
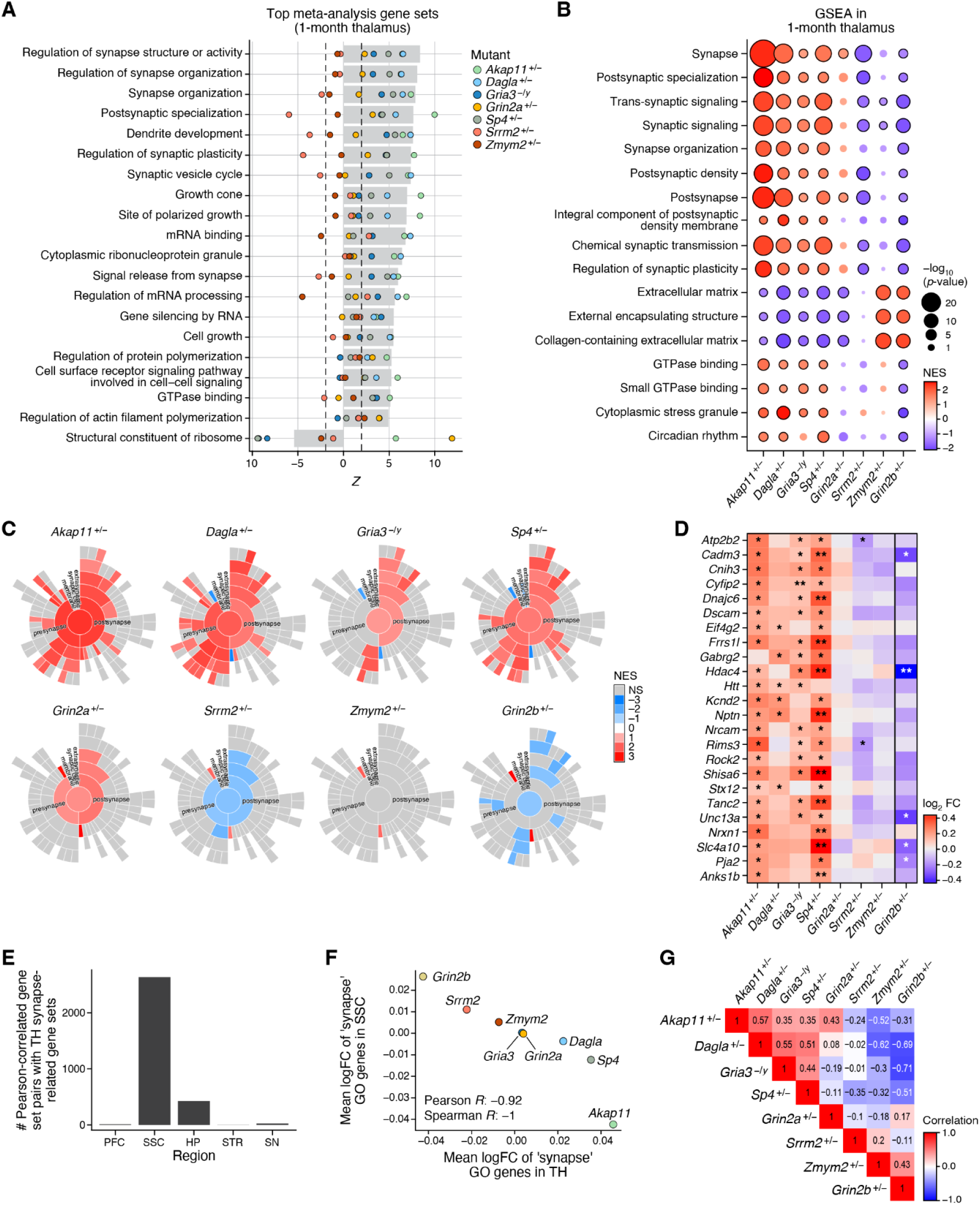
Transcriptomic changes in the thalamus show convergence among non-DD/ID associated SCHEMA mutants and divergence from DD/ID associated mutants. **A)** Top 20 most significant gene set changes by meta-analysis in 1 month thalamus across SCHEMA mutants. Gray bar represents Stouffer’s meta Z score, each point represents the individual gene set Z scores from each SCHEMA mutant. Values outside the dashed lines correspond to the Z score which represents nominal significance (p < 0.05). For the purpose of display, redundant gene sets were collapsed and only the strongest of the similar gene sets is plotted (see Methods) **B)** Individual GSEA results for gene sets which are convergent across the non-DD/ID mutants (FDR significant and directionally agreeing in all 4) as well as the circadian rhythm gene set. Color represents normalized enrichment score (NES), with red indicating upregulation and blue indicating downregulation. Bubble size represents significance, and black borders indicate FDR < 0.05 significance. **C)** Sunburst plots showing GSEA results using synGO gene sets. Color represents NES, and non significant terms (FDR > 0.05) are plotted in gray. See Supplemental Figure 5 for extended plot annotation, and full key to the sunburst plot can be found at the synGO portal https://www.syngoportal.org/ **D)** Log_2_ Fold Changes of the most consistently upregulated synapse genes across non-DD/ID mutants as determined by a rank-sum approach and meta-analysis. *: p < 0.05, **: FDR < 0.05. **E)** Number of significant (FDR < 0.05) gene set correlations between the NES of synapse related gene sets in 1 month thalamus and the NES of any gene set in other regions at 1 month across SCHEMA mutants, as determined by Pearson correlation. **F)** Mean log_2_FC of genes in the synapse GO gene set in thalamus and SSC for each mutant. **G)** Correlation of GSEA results across mutants. Spearman’s correlation for the union of nominally significant gene sets for each pair of mutants in 1 month thalamus. Number represents Spearman’s correlation coefficient.

These were largely driven by four of the SCHEMA mutants - *Akap11*, *Dagla*, *Gria3*, and *Sp4* (Figure 4A). Intriguingly, these are the four SCHEMA mutants in this study that are not genetically associated with DD/ID (hence termed “non-DD/ID-associated”) and which might therefore be considered more specific for SCZ/BP rather than general neurodevelopmental disorders. Besides the synapse-related gene sets, terms related to mRNA processing and binding (including cytoplasmic ribonucleoprotein granules, etc.; Figure 4A) were commonly altered in the meta-analysis.

Upon closer inspection of the synapse-related gene sets, we noted a striking divergence between the non-DD/ID-versus the DD/ID-associated SCHEMA mutants: namely, the synapse gene sets were upregulated in the non-DD/ID group (*Akap11*, *Dagla*, *Gria3*, *Sp4*), but unchanged or even downregulated in the DD/ID group (*Srrm2*, *Zmym2*) (Figure 4B-C). *Grin2a* does not fit neatly into either DD/ID or non-DD/ID category as certain missense mutations have been linked DD/ID, whereas protein-truncating/LoF mutations, as modeled here, have not ^5^.

Interestingly, the heterozygous *Grin2a* LoF mutants in our study had a thalamic transcriptomic phenotype that was intermediate between the above groups, though more similar to the non-DD/ID group (Figure 4B-C). Further, by utilizing synGO ontologies ^45^, the specific synapse-related gene set changes in thalamus were remarkably similar across these “non-DD/ID” SCHEMA mutants, with gene sets such as “presynaptic active zone”, “postsynaptic specialization”, and “postsynaptic density” upregulated in all four models (Figure 4C, Supplemental Table 9). At the individual gene level, the changes were also remarkably similar across the *Akap11*, *Dagla*, *Gria3*, and *Sp4* mutants, and the most commonly upregulated synapse-related genes (as identified by rank-sum analysis) in this group showed no change or were decreased in the DD/ID-associated mutants, with *Grin2a* again being intermediate (Figure 4D). Among the interesting synapse-related genes showing differential changes in DD/ID-versus non-DD/ID-associated SCHEMA mutants are *Atp2b2* (a calcium pump with emerging genetic association with bipolar/SCZ ^46^), *Gabrg2* (a GABA receptor subunit), and neurexin 1 (*Nrxn1*), deletions of which are associated with schizophrenia ^47^ (Figure 4D, Supplemental Table 2).

It is notable that based on transcriptomic phenotype in the thalamus, the *Grin2b* mutant, which is a genetic model of ASD/DD/ID, shows changes in synapse-related genes and gene sets that resemble those in *Srrm2* and *Zmym2* (DD/ID-associated SCHEMA genes) mutants but differ from those in *Akap11*, *Dagla*, *Gria3* and *Sp4* (non-DD/ID SCHEMA genes) mutants (Figure 4B, C, D).

We further investigated this divergence between DD/ID-associated and non-DD/ID-associated SCHEMA mutants in the thalamus by performing GSEA on disease-related gene sets (see Methods) derived from human GWAS and exome sequencing studies for various CNS disorders (Supplemental Figure 4A, Supplemental Table 10).

Interestingly, this analysis reinforced the separation of DD/ID-versus non-DD/ID mutants, based on enrichment scores for genes associated with several brain disorders, particularly schizophrenia (common-variant and SCHEMA genes), and neurodevelopmental delay (NDD) and ASD (genes identified from exome sequencing studies of ASD and NDD). Specifically, in these disease-associated gene sets, the strongest downregulation in thalamus was observed in *Grin2b* mutants, with weaker downregulation also seen in *Srrm2* and *Zmym2* mutants. By contrast, the non-DD/ID mutants (*Akap11*, *Dagla*, *Gria3*, *Sp4*) showed upregulation of these schizophrenia, NDD, and ASD gene sets in the thalamus (Supplemental Figure 4A).

Because the thalamus is an important sensory relay hub connected directly to multiple brain regions, we investigated whether the changes in synapse-related gene sets in the thalamus correlated with any gene set changes in other brain regions across mutants. We did this by calculating Pearson and Spearman correlations of normalized enrichment scores, (NES, which represent relative magnitude of enrichment) across mutants from GSEA (see Methods). The greatest number of significant gene set correlations with synapse-related gene sets in the thalamus was observed in the somatosensory cortex (SSC, Figure 4E, Supplemental Figure 4B, Supplemental Table 11). In fact, there was a consistent anti-correlation between synapse-related gene set enrichment in the thalamus and synapse-related gene set enrichment in the SSC - in mutants where synapse-related gene sets were upregulated in thalamus, they were downregulated or trending to downregulated in the SSC, and vice versa (Supplemental Figure 4C, Supplemental Table 11). Further, the average log_2_ fold change (log_2_FC) of the genes in the “synapse” gene ontology term shows a strong anticorrelation (Pearson’s R =-0.92, p = 0.001) between thalamus and SSC - mutants with a more positive average change in synapse genes in the thalamus have a more negative average change in the SSC and vice versa (Figure 4F). This strong degree of anti-correlation of altered gene expression in thalamus and SSC may be related to functional connectivity between the regions.

The divergence between DD/ID-associated versus non-DD/ID SCHEMA mutants in the thalamus extended beyond synapse-related terms. The overall correlation of GSEA results across the mutant models showed a clear pattern where *Akap11*, *Dagla*, *Gria3*, and *Sp4* were quite similar to each other and anti-correlated with *Srrm2*, *Zmym2*, and *Grin2b*, with *Grin2a* falling somewhere in the middle (Figure 4G). We searched for other gene sets that were significantly changed in the same direction in all of these non-DD/ID SCHEMA mutants, and we found that gene sets related to extracellular matrix were strongly downregulated and gene sets related to GTPase binding and stress granules were upregulated consistently across the non-DD/ID SCHEMA mutants (*Akap11*, *Dagla*, *Gria3*, *Sp4*) (Figure 4B). Remarkably, these same gene sets showed the opposite regulation or little change in the DD/ID-associated SCHEMA mutants (*Srrm2*, *Zmym2*), and consistent, significant opposite change in *Grin2b*. In this sense, the DD/ID-associated SCHEMA mutants showed overall similar GSEA directional changes in the thalamus as the mutant of *Grin2b*, a gene associated with DD/ID/ASD but *not* schizophrenia (Figure 4B, Supplemental Table 5).

Because the thalamus is involved in sleep regulation ^48^ and sleep disturbances are a common feature of schizophrenia ^49^, we additionally examined the “circadian rhythm” gene set and found that it also showed this divergence, with significant upregulation in the non-DD/ID SCHEMA mutants, a trend to downregulation in the DD/ID-associated SCHEMA mutants, and significant downregulation in the *Grin2b* mutant (Figure 4B).

## DISCUSSION

By systematic transcriptomic analysis of multiple mouse models of schizophrenia that are based on rare-variant human genetics (SCHEMA mutant mice), we have uncovered potential pathways of molecular dysregulation and systems-level disturbance that could be relevant to schizophrenia pathophysiology. We show that heterozygous (or hemizygous) mutations in a set of seven different SCHEMA genes (all of which substantially elevate risk of schizophrenia in humans) have quite heterogeneous effects on the brain transcriptome of mice. The transcriptome wide impact, the most affected brain region, the most affected age, and the precise genes and pathways that are altered, all showed considerable variation across the SCHEMA mutant models. This heterogeneity underscores that risk of schizophrenia–a diagnosis based solely on clinical symptoms rather than mechanistic understanding of disease–likely arises from many different causes, which can manifest with a variety of transcriptomic perturbations in the brain. In that context, it is unsurprising that different disease-associated genetic variants (such as these different SCHEMA mutations) do not alter the brain transcriptome in a uniform and drastic fashion. Rather, these SCHEMA mutations can have specific and different molecular effects across the brain, which is in keeping with the heterogeneity of schizophrenia as a psychiatric disorder.

Nonetheless, despite the heterogeneity of detailed transcriptomic phenotypes across mutants, we were able to identify convergences and overlaps among the SCHEMA mutants. Our systematic study – a coordinated characterization of multiple schizophrenia risk genes *in vivo* using non-hypothesis-driven transcriptomics across different brain regions and ages – has not been applied to such a large set of schizophrenia mouse models before. This comprehensive approach allows for the in-depth investigation of each individual SCHEMA gene when mutated in mice ^8,20–24^ as well as the meta-analyses of phenotypes across the different mutants, thereby helping to narrow down which effects are most likely to be relevant to the broader disease context. Previous studies have aimed to characterize multiple disease-associated genes with cell-specific mutations in ASD risk genes and identified convergent cellular mechanisms ^50,51^, but they were unable to tackle the systems-level question of which regions and networks are affected in the intact brain of multiple different mouse mutants.

A major conclusion from our study is that among the brain regions we surveyed, the striatum is the one that shows the most change and the strongest convergence across all of the SCHEMA mouse mutants characterized in this study. This conclusion, based on systematic, non-hypothesis-driven analysis, supports a long-standing concept that dysfunction of the striatum plays a key role in schizophrenia pathophysiology ^33^. Overall in the striatum, we found downregulation across SCHEMA mutants of genes and pathways related to synapses and mitochondrial respiration. One speculative interpretation of this transcriptomic change might be decreased neural activity in the striatum, however examining a set of activity-regulated genes – another potential readout of neuronal activity – revealed no consistent changes in striatum across mutants. Single-cell transcriptomic analysis may therefore be informative to better infer changes in activity of neurons in the striatum.

In our meta-analysis we also identified a high impact and a convergence of transcriptomic changes in the thalamus, particularly among SCHEMA genes that do not have an association with DD/ID (*Akap11*, *Dagla*, *Gria3*, *Sp4*). Although less well-studied than PFC and striatum, there is growing evidence of thalamic involvement in schizophrenia ^52–54^. This convergent transcriptomic phenotype in the thalamus among non-DD/ID SCHEMA genes, and to a lesser extent *Grin2a*, may point to thalamic disturbances that are more specific to schizophrenia or psychosis spectrum disorders (*Sp4* and *Akap11* are also associated with bipolar disorder) than with severe neurodevelopmental disorders such as DD/ID/ASD. There were many gene sets that followed this divergent pattern between “SCZ/BP-specific” and DD/ID-associated SCHEMA mutants, but most strikingly, we found the upregulation of synapse-related gene sets in the thalamus to distinguish the non-DD/ID SCHEMA mutants from the DD/ID-associated SCHEMA mutants and *Grin2b*. It is also intriguing that this change in thalamus is anti-correlated with alterations in synapse-related gene sets in the somatosensory cortex - potentially insinuating altered functional connectivity or aberrant communication in the sensory processing pathway from thalamus to sensory cortex. This is notable because functional MRI studies in human schizophrenia patients have found thalamic over-connectivity with bilateral sensory-motor cortices ^55^, and further have identified correlation between thalamic connectivity and cognitive deficits and negative symptoms ^56^. In this study we do not have data from other neocortical areas (except PFC), but exploration of additional cortical areas is an interesting direction for follow-up. The anti-correlation of synapse-related gene changes between the thalamus and SSC also occurs strongly in the ASD-linked *Grin2b* mutant, but in the opposite direction; in that context, we note that altered sensory processing is increasingly recognized as a characteristic feature of autism ^57^.

In terms of detailed circuitry and specific cell types, this study is limited in resolution due to the analysis of bulk RNA-seq data. Our strategy, however, allowed us to broadly survey many brain regions at two different ages, allowing us to investigate how widespread brain regions and systems are differentially impacted during CNS maturation from juvenile to young adult stages. In its favor, bulk RNA-seq does address total cellular RNA as opposed to just the small fraction of transcripts present in the nucleus. Future assessment of these brain regions of interest using the more costly single-nucleus or single-cell RNA-seq should enable a deeper interpretation of the convergent changes we identified at the bulk level. Spatial transcriptomics is another modality that would be highly informative to better understand the dysfunction in SCHEMA mutant mice. Even with bulk RNA-seq, it is clear that age has a substantial influence on the transcriptomic phenotype of all the SCHEMA mutants across every brain region examined, both in terms of transcriptomic impact and nature and direction of genes and pathways affected. One of many examples of this age-dependence is illustrated by the *Grin2a* mutant, which shows opposite direction of enrichment of human schizophrenia DEGs at 1 month versus 3 months of age. Overall our observations underscore the importance of age and stage of brain maturation, and the large difference between various brain regions, when considering the impact of genotype on the brain’s gene expression patterns.

This study highlights the usefulness of simultaneous and systematic evaluation of many different genetic models of a given complex disease, rather than just a single example one at a time. We have shown that like schizophrenia itself, its genetic risk factors are very heterogeneous and their heterozygous loss of function mutation can lead to a wide variety of molecular changes in the brain. Despite this overall heterogeneity, however, we were able to identify interesting patterns of transcriptomic change that are common across all SCHEMA mutants or shared within defined subsets of SCHEMA mutants. Deeper investigation of these convergent transcriptional phenotypes–as well as extension to additional SCHEMA and non-SCHEMA mutant mice– could help us to understand the shared and the disparate mechanisms that give rise to the clinical syndrome of schizophrenia and to formulate new therapeutic approaches.

## METHODS

### Mouse models

*Grin2a^-/-^* mice ^58^ were obtained from the RIKEN BioResource Research Center (RBRC02256) and were crossed with wild-type C57/BL6J mice (Jackson Laboratory, #000664) to generate *Grin2a^+/-^* mice. Mice were bred and maintained as previously described ^8^.

*Akap11^+/-^* heterozygous mice (B6.Cg-Akap11[tm1.2Jsco/J]) ^59^, were obtained from The Jackson Laboratory following cryorecovery (strain #028922). Mice were bred and maintained as previously described ^22^.

*Zmym2*^+/-^ (C57BL/6NJ-Zmym2 em1(IMPC)J/Mmjax) mice were obtained from The Jackson Laboratory. Mice were bred and maintained as previously described ^24^.

The *Srrm2 ^em1(IMPC)Bay^* mouse line was used for this study. Mutant mice were re-derived from cryopreserved germplasm by in-vitro fertilization at Harvard Genome Modification Facility (Cambridge, MA), and back-crossed with C57BL/6J (The Jackson Laboratory, #000664).

Heterozygous mice and wild-type littermates were bred and maintained as previously described23.

*Sp4* mutant mice were generated using the CRISPR/Cas9 technique to have an early terminating stop codon in exon 3 as described in Kwon et al., 2024. Mice were bred and maintained as previously described ^21^.

*Gria3* mutant mice were generated using a double-stranded DNA donor knock-in strategy to introduce the GRIA3R210X mutation as described in Huang et al., 2024 ^20^. Mice were bred and maintained as previously described^20^.

Frozen whole brain tissues of *Dagla* mutant mice and their wild-type littermates, as previously described ^60^, were kindly provided by Dr. Aron Lichtman (Virginia Commonwealth University).

Frozen whole brain tissues of *Grin2b^+/C456Y^* and their wild-type littermates, all on the genetic background of C57BL/6J, were kindly provided by Dr. Eunjoon Kim (Department of Biological Sciences, Korea Advanced Institute of Science and Technology, South Korea).

Brain dissection was performed as previously described ^8,20–24^. For clarity, the PFC represents a dissection of the medial prefrontal cortex, the hippocampus represents a dissection of the dorsal hippocampus, and the striatum represents a biopsy punch of the dorsal striatum.

The thalamus and substantia nigra were collected with a biopsy punch, the somatosensory cortex was dissected by hand.

### RNA-sequencing

RNA-sequencing was performed as previously described ^8,20–24^. Briefly, RNA was isolated from micro-dissected tissues using the RNeasy Mini Kit (Qiagen) following the manufacturer’s instructions. Bulk sequencing libraries were prepared using a TruSeq Stranded mRNA Kit (Illumina) following the manufacturer’s instructions. A 10 nM normalized library was pooled, and sequencing was performed on a NovaSeq S2 (Illumina) with 50 bases each for reads 1 and 2 and 8 bases each for index reads 1 and 2.

### RNA-seq data processing

Raw FASTQ files from each sequencing experiment were aligned to a standardized reference (genome FASTA file and transcriptome GTF were extracted from the CellRanger mm10 reference and used for alignment and quantification) using a Nextflow V1 pipeline ^61^ (https://github.com/seanken/BulkIsoform/blob/main/Pipeline). Data that were not previously published, but generated in the same study ^8^, were included for *Grin2a* and *Grin2b* mutant substantia nigra, and *Grin2a* wild-type 3 month PFC. Data from the 3 month substantia nigra for *Srrm2* mutants were not analyzed due to low number of replicates. Quantification was performed by Salmon ^62^ (version 1.7.0, with arguments-l A--posBias--seqBias –gcBias--validateMapping), using a Salmon reference built with the salmon index command with genomic decoys included. Quality metrics were obtained from STAR alignment (version 2.7.10a) ^63^ and Picard tools CollectRnaSeqMetric command ^64^ (version 2.26.7)(Supplemental Table 1).

Samples with poor quality were removed from analysis (Supplemental Table 1).

### Differential Expression

Differential expression (DE) analysis was performed between heterozygous mutant and wild-type mice for each individual experiment (SCHEMA mutant, age, and brain region) using the Wald test in R package DESeq2 ^65^ (version 1.34). Salmon output was loaded using R package tximport ^66^ (version 1.22). All genes that had at least 10 counts across all samples in each experiment were included in our analysis. Due to evidence of dissection variability in striatum by excitatory markers, DESeq2’s PC1 was added as a covariate for differential expression in 1 month *Akap11* and 1 month *Grin2b*. Log_2_ fold change shrinkage was applied using the “normal” shrinkage estimator in DESeq2. DESeq2’s default results filtering was used and all genes in a given experiment with padj = NA were removed from downstream analysis. For *Akap11* 1 month hippocampus and thalamus, and *Grin2a* 1 month hippocampus, the default filtering removed the majority of genes, so we changed the filtering for the heterozygous vs. WT comparison to match the less aggressive filtering in the knockout vs. WT comparison.

### Gene Set Enrichment Analysis

Gene set enrichment analysis (GSEA) was performed using R package fgsea ^67^ (version 1.20). GSEA was run separately for each set of DE results (i.e., each mutant, brain region, and age), using DESeq2’s “stat” column to rank the genes. GSEA for meta-analysis was performed using the GO database with the recommended minimum gene set size of 15 and maximum gene set size of 500, and the GO gene sets were downloaded using R package AnnotationDbi ^68^ (version 1.56.2). For plotting purposes only, GSEA was also run with a maximum gene set size of 2500 so that gene sets of interest (e.g. “synapse”) could be examined. GSEA was also run separately on the synGO database ^45^ (version 1.1), and in the cases where the mutant gene was in a synGO gene set, it was removed from the input. For comparison to human genetic data from psychiatric and neurological diseases, GSEA was run on curated gene sets as previously described ^8^. Finally, for comparison to human PFC RNA-seq data, gene sets were created for schizophrenia upregulated (padj < 0.05 & log_2_FC > 0) and downregulated (padj < 0.05 & log_2_FC < 0) genes in BrainSeq Phase II ^44^ and GSEA was run using these gene sets. FDR was calculated across all experiments for this specific human-SCZ DEG gene set analysis.

### TRADE

For each mutant, we re-estimated differential expression effects using DESeq2 without shrinkage. We included surrogate variables estimated with SVA ^69^ as fixed effects in the DESeq2 model. We picked the number of surrogate variables using the permutation-based’be’ method included in SVA.

We then submitted the DESeq2 summary statistics for analysis with TRADE ^29^ to estimate the distribution of differential expression effects. Briefly, TRADE fits a flexible mixture distribution to the gene-wise log_2_FCs and standard errors to estimate the distribution of effect sizes across genes. The key output of the TRADE analysis is the “transcriptome-wide impact”, the variance of the effect size distribution, which is a measure of the overall impact of a perturbation on the transcriptome.

To assess significance of these estimates, we repeated the above analysis 1000 times with permuted genotype labels, and computed the proportion of permutation transcriptome-wide impact estimates that were larger than the empirical estimate.

To meta-analyze transcriptome-wide impact across experiments, we used Fisher’s method to combine p values from all SCHEMA mutants into a meta p value.

To obtain meta-analytic transcriptome-wide impact point estimates and standard errors for visualization, we used a heuristic approach. We used point estimates and permutation p values to estimate standard errors using the quantile function of the standard normal distribution, i.e., assuming that the sampling distribution of the estimate is normal, which is asymptotically true but may not hold at finite sample size. We note that this heuristic assumption does not affect the validity of the estimated meta p values. We used precision-weighted fixed effects meta-analysis to obtain meta point estimates and standard errors from the experiment-wise point estimates and standard errors.

### Meta-analysis

Stouffer’s weighted Z-score meta-analysis was performed across all SCHEMA mutants separately for each age and brain region to identify the strongest and most consistent changes across mutants. For each gene set in GSEA or gene in DE, the individual Z score (Z_i_) for each mutant was calculated by applying R’s qnorm function to the p-value divided by 2, then multiplying by the sign of the NES or log_2_FC and-1. Then, the meta-analysis statistic (Z) was calculated with the following formula:

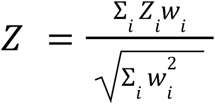

Weights (w_i_) were defined as the square root of the total N (mutant + wild type) for each experiment. The meta-Z score was then converted to a p-value and FDR correction was applied within each brain region and age. When counting the number of significant genes by meta-analysis, the mutant genes were not included. For plotting purposes only, meta-analysis results were filtered to reduce redundancy in GO terms using the function go_reduce from R package RHReynolds/rutils ^70^ (version 0.99.2). We input the absolute value of Stouffer’s Z as the score for each GO term, and the threshold was set to 0.9 for striatum and thalamus, and 0.7 for PFC.

### Correlation

For cross-mutant correlation plots, Spearman correlation was performed on the NES from GSEA for the union of nominally significant (p < 0.05) gene sets for each pair of mutant models. For correlation with human RNA-seq data, Spearman correlation was performed on the logFCs of the union of nominally significant genes between the PFC DE results from each mutant at each age and the human PFC differential expression results ^44^.

For correlation between thalamus synaptic gene sets and gene sets in all other regions, in 1 month thalamus, GO gene sets which contained “synap”, “dendrit”, and “axon” and were nominally significant in at least 3 mutants were selected. For each other brain region, all gene sets that were nominally significant in at least 3 mutants were selected. Then, both Spearman and Pearson correlations were performed on the NES across SCHEMA mutants for each gene set. P-values for each correlation were derived from a t-statistic calculated as:

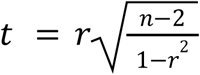

The p-values were then FDR-corrected to determine significance.

## Data availability

Data for *Akap11* ^22^, *Gria3* ^20^, *Grin2a* ^8^, *Sp4* ^21^, *Srrm2* ^23^, *Zmym2* ^24^, and *Grin2b* ^8^ mutants have been previously described. Data for *Dagla* mutants and substantia nigra data from *Grin2a* and *Grin2b* mutants will be uploaded to GEO prior to publication. Extended results data are available upon request.

## Code availability

Code will be available in a GitHub repository at https://github.com/StanleyCenter-ShengLab/PerzelMandell_etal_2024.

## Acknowledgements

We thank Kris Dickson and members of the Sheng and Levin Labs for valuable input, Leslie Gaffney for assistance in improving and assembling figures, Eunjoon Kim for sharing *Grin2b* brain tissues, and Calwing Liao for insights into meta-analysis methods.

## Competing interests

MS is cofounder and scientific advisory board (SAB) member of Neumora Therapeutics and serves on the SAB of Biogen, Proximity Therapeutics, and Illimis Therapeutics.

## Contributions

KPM and MS conceived and planned the work. KPM performed data processing, analysis, and interpretation, with guidance from SKS and supervision from JZL and MS. KPM wrote the manuscript and MS and JZL substantially revised it. ZF, WH, SA, MJK, BS, KB, NS, and AAN generated and shared the RNA-sequencing data. SKS and AN developed and provided code for data processing and analysis. AHL and LOB provided tissues for data generation.

Supplemental Table 1: Samples and QC

Supplemental Table 2: Differential Gene Expression Results Supplemental Table 3: TRADE results

Supplemental Table 4: Convergence results

Supplemental Table 5: GO Gene Set Enrichment Analysis Results Supplemental Table 6: GSEA Stouffer’s Meta-Analysis Results

Supplemental Table 7: Differential Expression Stouffer’s Meta-Analysis Results Supplemental Table 8: GSEA using DEGs in human SCZ PFC

Supplemental Table 9: synGO GSEA Results

Supplemental Table 10: Disease Gene Set GSEA in 1 month thalamus

Supplemental Table 11: Correlation between 1 month thalamus GSEA and other brain regions

