## Supplemental Figures for "Meta-analysis of the brain transcriptomes of multiple genetic mouse models of schizophrenia highlights dysregulation in striatum and thalamus"

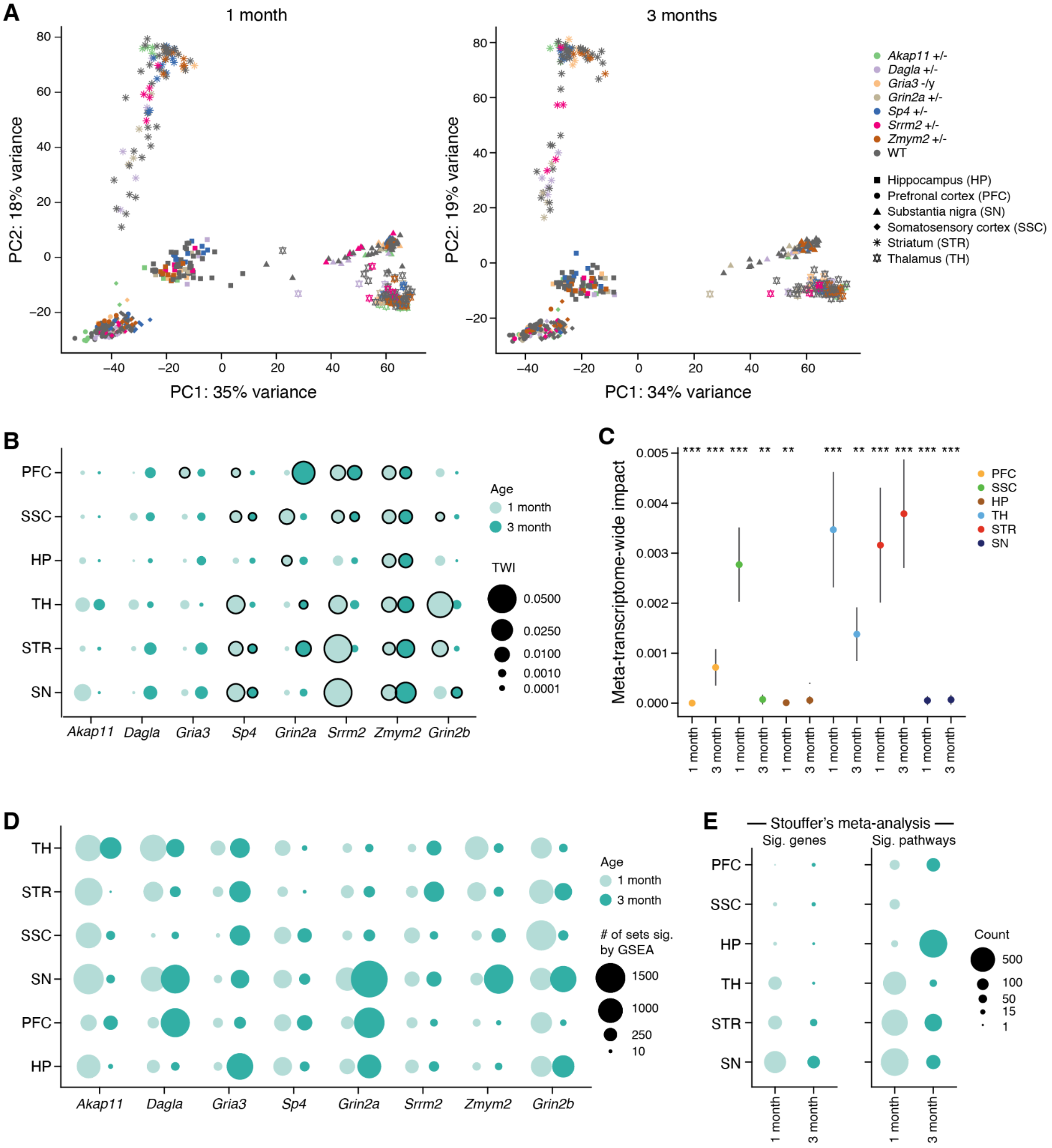


**Supplemental Figure 1. Breadth of RNA-sequencing and transcriptomic impact of SCHEMA mutants**

**A)** PCA of all samples included in this meta-analysis, with color representing genotype and shape representing brain region **B)** Transcriptome-wide impact scores for each mutant, brain region, and age, as indicated. Bubble size indicates TWI score, color indicates age, and a black ring indicates nominal significance ($p < 0.05$). **C)** Meta-TWI as meta-analyzed across SCHEMA mutants for each age and brain region. **D)** Number of significant gene sets by GSEA for each mutant, brain region, and age. **E)** Number of significant genes and GO gene sets (pathways) by Stouffer's meta-analysis across SCHEMA mutants for each brain region and age.

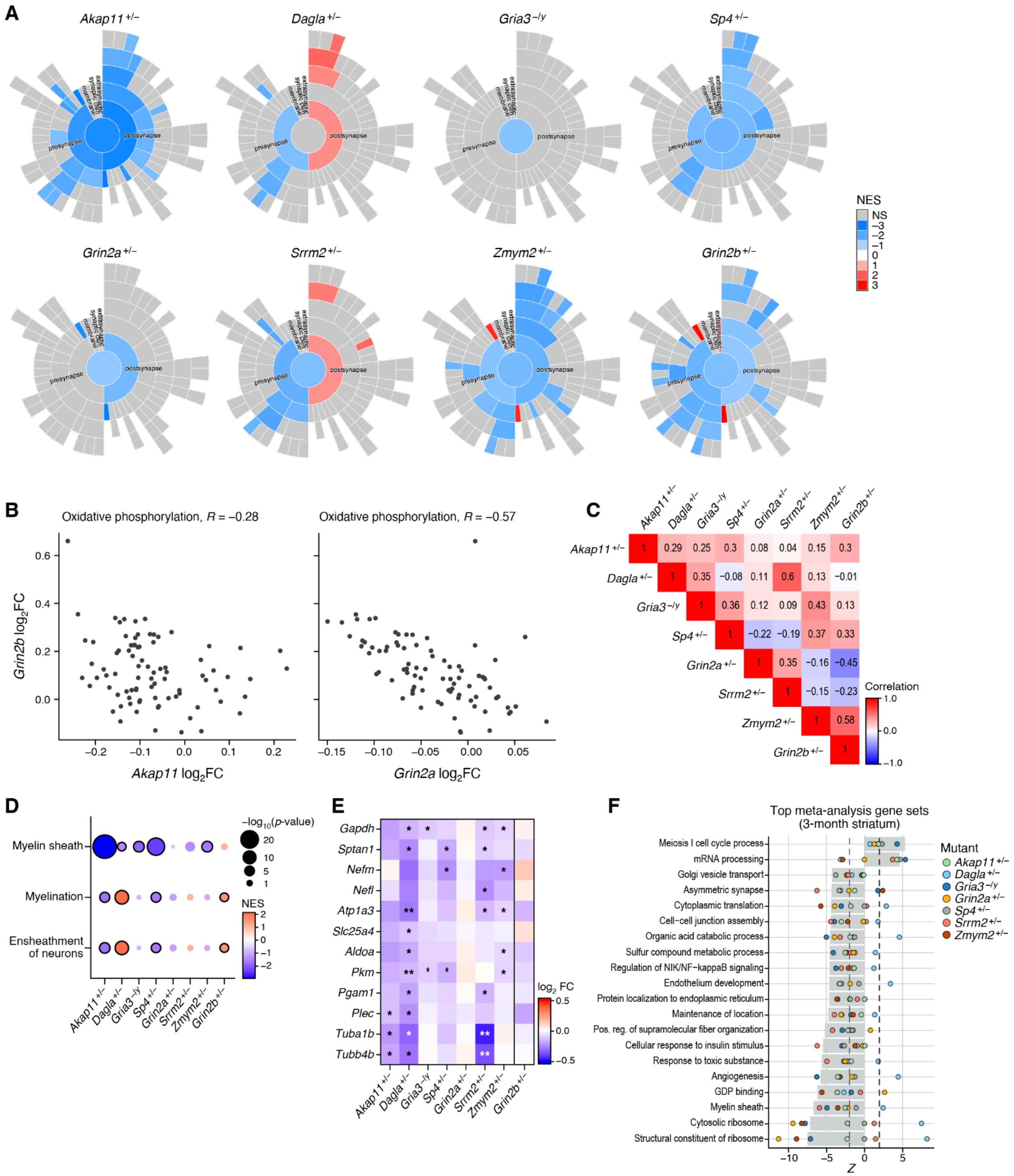


**Supplemental Figure 2. Convergence of pathway-level transcriptomic changes in SCHEMA mouse striatum**

**A)** SynGO sunburst plots for GSEA in 1 month striatum. Color represents NES, and non significant terms (FDR > 0.05) are plotted in gray. See Supplemental Figure 5 for extended plot annotation, and full key to the sunburst plot can be found at the synGO portal https://www.syngoportal.org/ **B)** Correlation of $log_2FC$ for all genes in the "oxidative phosphorylation" GO gene set between SCHEMA mutants (Grin2a and Akap11) and Grin2b **C)** Correlation of GSEA results across mutants. Spearman's correlation was performed on the union of nominally significant gene sets for each pair of mutants in 1 month striatum. Number represents Spearman's correlation coefficient. **D)** Individual GSEA results for myelin related gene sets in 1 month striatum. **E)** Most consistently downregulated genes in the myelin sheath gene set across SCHEMA mutants in 1 month striatum as determined by a rank-sum approach. * : $p < 0.05$, ** : FDR < 0.05. **F)** Top 20 most significant gene set changes by meta-analysis in 3 month striatum across SCHEMA mutants. Gray bar represents Stouffer's meta Z score, each point represents the individual gene set Z scores from each SCHEMA mutants. Dashed lines correspond to the Z score which represents nominal significance ($p < 0.05$). For the purpose of display, redundant gene sets were collapsed and only the strongest of the similar gene sets is plotted (see Methods)

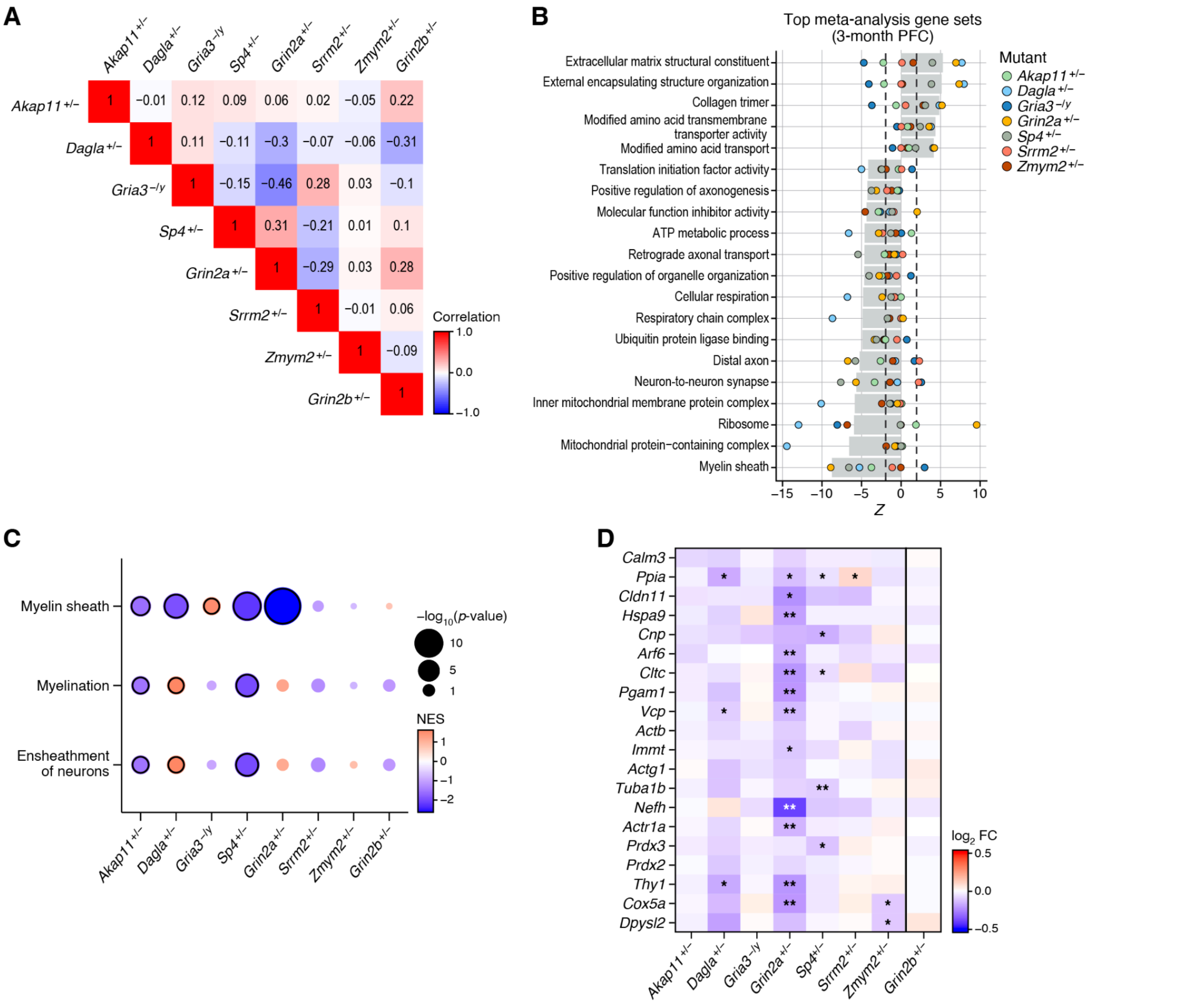


**Supplemental Figure 3. Heterogeneity of transcriptomic change in SCHEMA mouse PFC**
**A)** Correlation of DE results in PFC across mutants. Spearman's correlation was performed on the union of nominally significant genes for each pair of mutants in 1 month PFC. Number represents Spearman's correlation coefficient. **B)** Top 20 most significant gene set changes by meta-analysis in 3 month PFC across SCHEMA mutants. Gray bar represents Stouffer's meta Z score, each point represents the individual gene set Z scores from each SCHEMA mutants. Dashed lines correspond to the Z score which represents nominal significance ($p < 0.05$). For the purpose of display, redundant gene sets were collapsed and only the strongest of the similar gene sets is plotted (see Methods) **C)** Individual GSEA results for myelin related gene sets in 3 month PFC. **D)** Most consistently downregulated genes in the myelin sheath gene set across SCHEMA mutants in 3 month PFC as determined by a rank-sum approach. * : $p < 0.05$, ** : FDR < 0.05

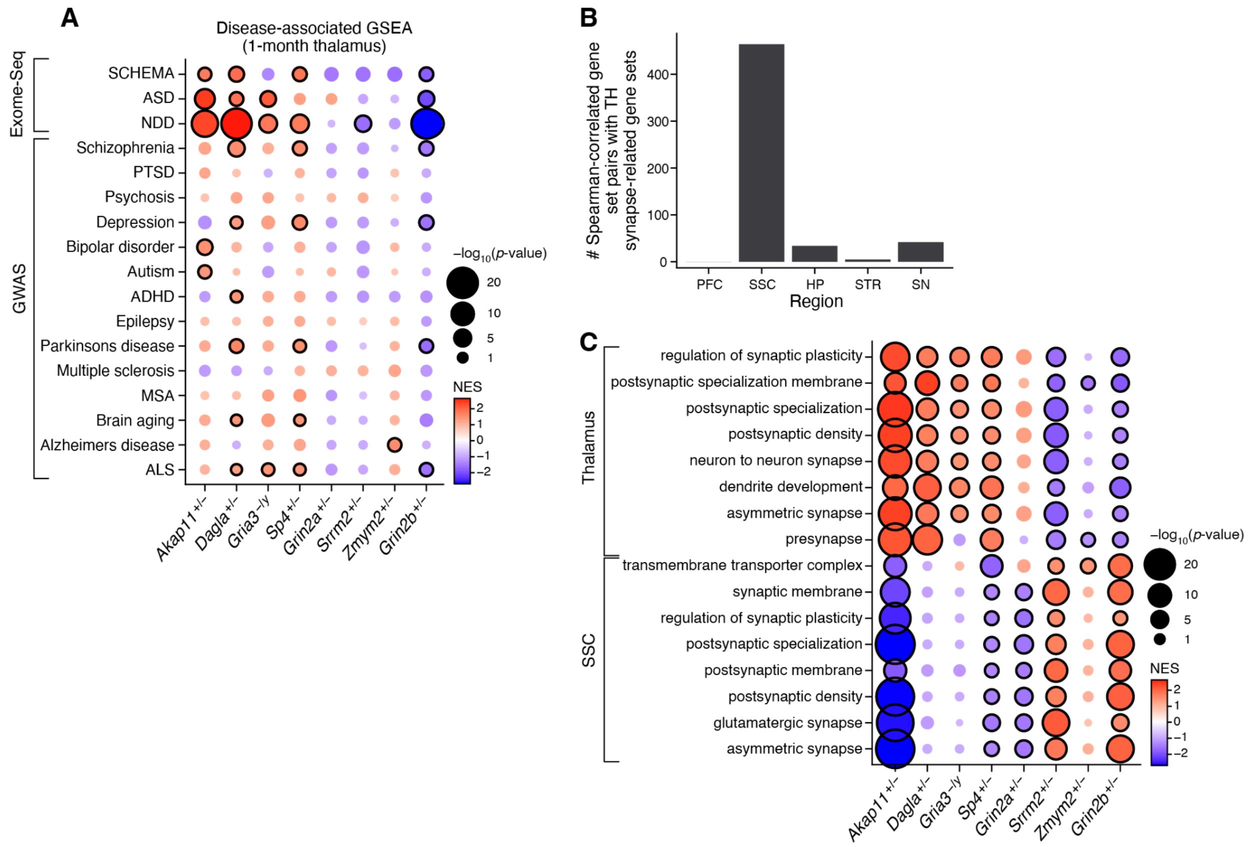


**Supplemental Figure 4. Divergence between DD/ID- and non-DD/ID associated SCHEMA mutants in synaptic disease-related pathways**

**A)** GSEA results in 1 month thalamus from disease gene sets based on GWAS and exome sequencing studies. **B)** Number of significant (FDR < 0.05) gene set correlations between the NES of synapse related gene sets in 1 month thalamus and the NES of any gene set in other regions at 1 month across SCHEMA mutants, as determined by Spearman correlation. **C)** GSEA results from gene sets involved in top synapse-related gene set anticorrelations in 1 month thalamus and SSC.

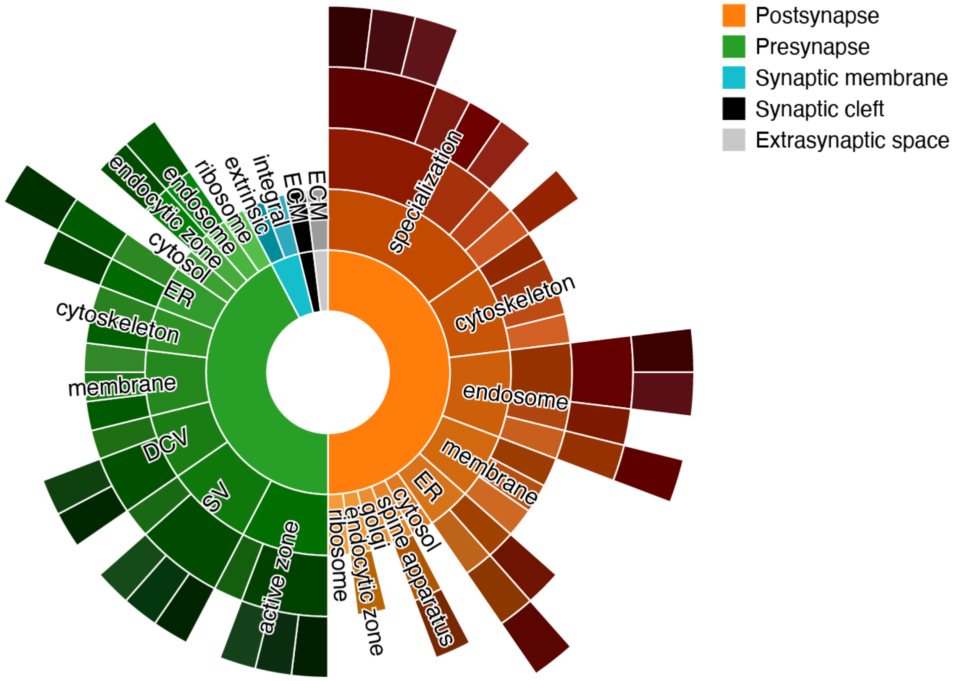


**Supplemental Figure 5. Extended annotation of synGO cellular component sunburst plot.** Annotation of all plot segments can be found at https://www.syngoportal.org/.

Supplemental Table 1: Samples and QC
Supplemental Table 2: Differential Gene Expression Results
Supplemental Table 3: TRADE results
Supplemental Table 4: Convergence results
Supplemental Table 5: GO Gene Set Enrichment Analysis Results
Supplemental Table 6: GSEA Stouffer's Meta-Analysis Results
Supplemental Table 7: Differential Expression Stouffer's Meta-Analysis Results
Supplemental Table 8: GSEA using DEGs in human SCZ PFC
Supplemental Table 9: synGO GSEA Results
Supplemental Table 10: Disease Gene Set GSEA in 1 month thalamus
Supplemental Table 11: Correlation between 1 month thalamus GSEA and other brain regions
